# Efficient Formation of Single-copy Human Artificial Chromosomes

**DOI:** 10.1101/2023.06.30.547284

**Authors:** Craig W. Gambogi, Elie Mer, David M. Brown, George Yankson, Janardan N. Gavade, Glennis A. Logsdon, Patrick Heun, John I. Glass, Ben E. Black

## Abstract

Large DNA assembly methodologies underlie milestone achievements in synthetic prokaryotic and budding yeast chromosomes. While budding yeast control chromosome inheritance through ∼125 bp DNA sequence-defined centromeres, mammals and many other eukaryotes use large, epigenetic centromeres. Harnessing centromere epigenetics permits human artificial chromosome (HAC) formation but is not sufficient to avoid rampant multimerization of the initial DNA molecule upon introduction to cells. Here, we describe an approach that efficiently forms single-copy HACs. It employs a ∼750 kb construct that is sufficiently large to house the distinct chromatin types present at the inner and outer centromere, obviating the need to multimerize. Delivery to mammalian cells is streamlined by employing yeast spheroplast fusion. These developments permit faithful chromosome engineering in the context of metazoan cells.

**One-Sentence Summary:** A quarter century after the first human artificial chromosomes, a solution to their uncontrolled multimerization is achieved.

---

Yeast artificial chromosomes (YACs) (*1–3*) are typically 0.1-1 Mbp and permitted triumphs of molecular biology including the cloning of large disease genes (*4*) and the generation of entire synthetic prokaryotic genomes (*5*, *6*). They also provided the foundation for the generation of entirely synthetic budding yeast chromosomes (*7*). ‘Writing’ new chromosomes, or even entire genomes, is an aspiration for synthetic biologists working in diverse eukaryotes, including in mammalian and plant systems (*8*, *9*), because it would enable applications of genome engineering across research, biotechnology, and health-related landscapes (*8*). For instance, one could engineer cancer resistance into new therapeutic cell lines.

Human artificial chromosomes (HACs) were developed ∼25 years ago (*10–12*) and are typically 1-10 Mbp in their functional form after their establishment in cells. They potentially paved the way for their deployment for applications in many eukaryotic systems where a specific key chromosomal locus, the centromere, is typically more than a thousand times larger than a budding yeast point centromere and is functionally defined not by a particular sequence but by an array of nucleosomes containing a histone H3 variant, CENP-A (*13*). Unlike YACs, *de novo* formation of HACs has obligatorily involved multimerization of the initially input DNA construct (typically 100-200 kb bacterial artificial chromosomes [BACs]), creating functional HACs with a variable number of multimers (typically >40-fold) (*14–16*). The multimerization and the uncontrolled rearrangement of the input DNA that accompanies it during the early steps of HAC formation has severely hindered their development towards their broader promise for synthetic biology and therapeutic applications (*17*). We have now overhauled the design and delivery of HACs: instead of trying to optimize the multimerization process, we sought to bypass it completely. Here we report success in forming single-copy HACs at an overall efficiency of *de novo* establishment that surpasses all earlier versions.

## Results

### An overhauled platform for efficient HAC formation

We predicted that to remain single-copy and avoid multimerization the initial construct would need to be larger than the BAC-based HAC constructs of earlier versions (*14–16*). This is based on the understanding that centromeres requires multiple domains with distinct functions that are spatially separated at mitosis when cohered sister chromatids align on the microtubule-based spindle (*18*). While the centromeric region harboring CENP-A nucleosomes that participates in assembling the mitotic kinetochore typically discontinuously spans ∼75-300 kb (*19–22*), the inner centromere is another largely heterochromatic region that regulates sister chromatid cohesion and a quality control mechanism (termed “error correction”) that monitors bipolar spindle attachment. We reasoned that BAC-based HAC constructs, which typically start in the 100-300 kb size-range (*14–16*), can likely only form when multimerization occurs because they must achieve the larger size required to accommodate formation of both distinct chromatin domains that define a functional centromere. Conversely, we reasoned that starting with a larger initial construct will bypass this requirement, allowing HACs to form more frequently and without multimerization.

To test our prediction, we devised a scheme that employs three recent technical advances to build and test a single-copy HAC construct (Fig. S1A). First, YAC constructs are readily generated in the 0.5-2 Mb size range (*5*, *23*) through transformation-associated recombination (TAR) cloning (*24*). Second, bypassing the requirement for long (>40 kb) stretches of highly repetitive centromere DNA (α-satellite) for HAC formation (*16*) permits the use of non-repetitive DNA. This is conducive to TAR cloning because it is not compatible with long repetitive sequences (*25*). Third, large YAC constructs can be efficiently delivered to mammalian cells via optimized fusion with yeast spheroplasts (*23*), potentially leading to a marked increase in independent HAC formation events relative to what has been achieved with low-efficiency transfection-based delivery of BAC-based HAC vectors in prior versions (*14–16*).

The HAC template was constructed through TAR assembly starting with a YAC harboring 550 kb of *M. mycoides* genomic DNA (*6*), 4q21 BAC^LacO^ (*16*), and linkers for recombination that also include a yeast auxotrophic marker and a mammalian expression cassette for mCherry (Fig. S1A). *M. mycoides* genomic DNA was chosen because it represents a heterologous DNA sequence that is known to be readily propagated in budding yeast (*6*). It serves as a non-coding sequence in the context of a eukaryotic cell and is not expected to elicit unintended or detrimental impact on HAC formation or cell function. Further, *M. mycoides* DNA has already been efficiently delivered to cultured human cells (*23*) and because it is a unique non-human DNA sequence it allows for unambiguous detection of HACs. 4q21 BAC^LacO^ was chosen because it is the only HAC construct comprised of non-repetitive DNA that has been demonstrated to form functional HACs, instead of the 40-200 kb of highly repetitive α-satellite-based BAC constructs that prior HAC studies have used (*14–16*, *26*). We termed the new construct, YAC-*Mm*-4q21^lacO^.

For recipient cells, we used the HT1080^Dox-inducible^ ^mCherry-LacI-HJURP^ line in which 4q21 BAC^LacO^ based HACs were seeded with CENP-A nucleosomes (*16*). The HT1080 background, in general, was chosen because it is the one in which HAC formation has historically been performed (*11*, *12*, *14–16*, *26*) due to its chromatin state that is permissive to occasional centromere formation (*27*). We also generated a second recipient line, U2OS^Dox-inducible^ ^mCherry-LacI-HJURP^, since the U2OS background are established as an efficient recipient of YACs via spheroplast fusion (*23*). Both of our chosen recipient lines were first optimized for spheroplast fusion conditions (Fig. S2) and then subjected to HAC formation assays with YAC-*Mm*-4q21^lacO^ (Figs. 1 and S2). Following spheroplast fusion, we noted that, unlike prior HAC assays where only ∼40 surviving colonies emerge in 2-3 weeks, a nearly confluent monolayer of G418-S-resistant cells was present after 8 days of selection. For both recipient cell types, a substantial proportion (42 +/-9% and 46 +/-5%) of the neomycin-resistant cells harbor HACs. Most or all of these are substantially smaller in size (<1 μm)(Fig. 1B-D) than the multimerized HACs of prior generations (∼2 μm)(*14–16*). Without induction of mCherry-LacI-HJURP, there was only a very small proportion of HACs, with the majority of cells with detectable FISH signal coming from an integration into a natural chromosome of the recipient cell (Fig. 1C). Our initial findings, therefore, strongly indicate exceedingly high efficiency of YAC delivery, robust HAC formation rates upon seeding CENP-A nucleosome assembly, essentially uniform avoidance of any or all of the high levels of multimerization that have accompanied prior systems for *de novo* HAC formation, and no restriction to the specific cell line (HT1080) to which prior generations of HACs were confined.

**Figure 1:**
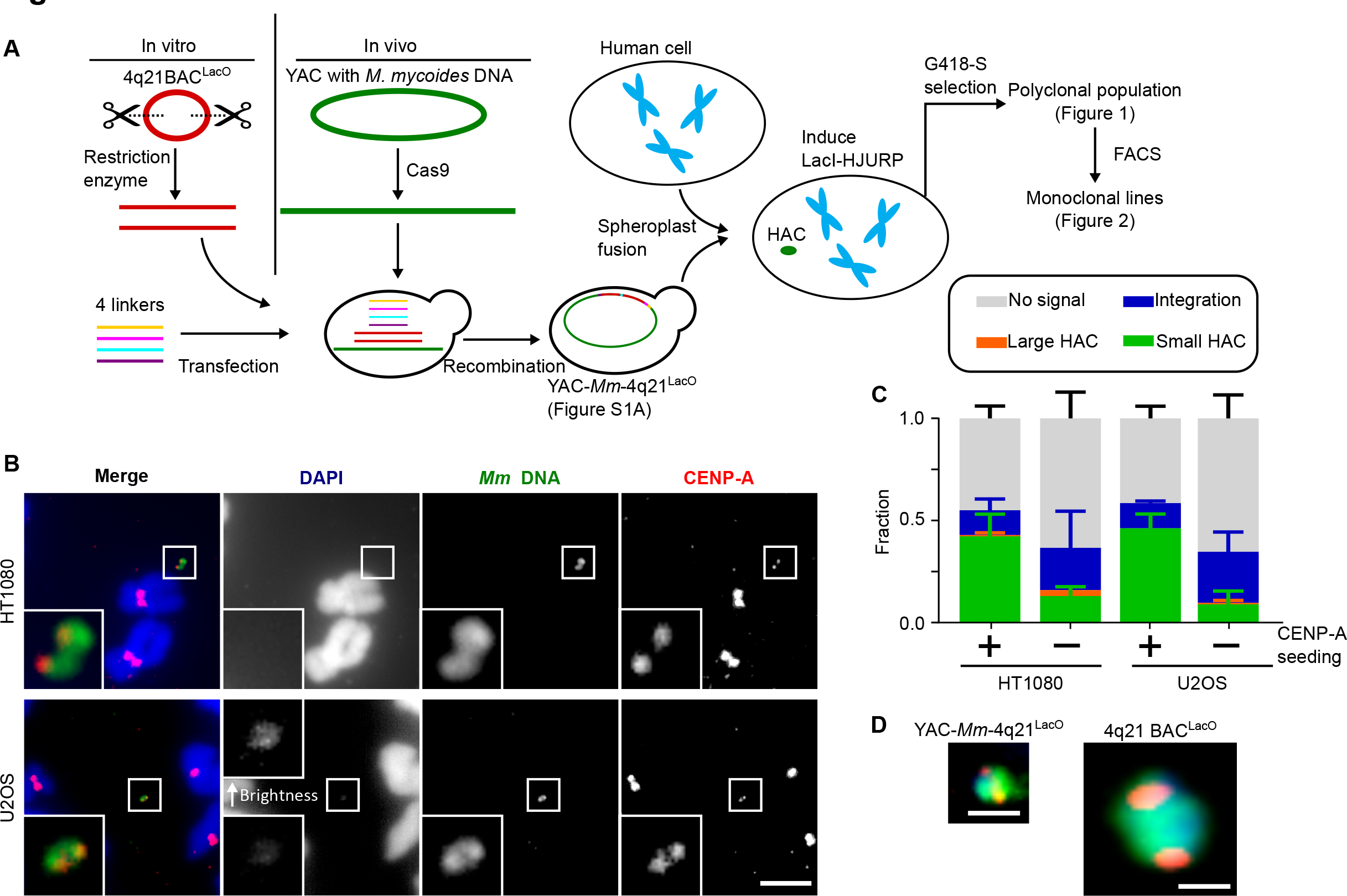
760 kb HAC constructs efficiently acquire centromeres and exist as autonomous chromosomes. A) Schematic of approach to generate a HAC. B) Representative images of a single-copy HAC generated in HT1080 and U2OS cells. Insets: 5x magnification. Bar, 5 μm. See also Table S1. C) Quantification of proportion of “small HACs” (FISH signal spans less than 1 μm), “large HACs” (FISH signal spans greater than 1 μm), “integrations” and “no signal” spreads generated from HAC formation assays. The mean (+/-SD) is shown. D) Comparison of size of a HAC made from YAC-*Mm*-4q21^LacO^ and a multimerized HAC made from 4q21 BAC^LacO^. Both HACs are shown at the same scale. Bar, 1 μm.

### YAC-*Mm*-4q21^LacO^ HACs Harbor Multi-domain Centromeres for Faithful Inheritance

We next sought to test the degree to which the centromeres on the HACs could support mitotic function. Starting from the polyclonal population of cells that survived G418-S selection (Fig. 1), we isolated eight monoclonal cell lines harboring HACs and measured the proportion of cells in each harboring a detectable HAC (Fig. 2A,B). Overall, the majority of cells within a clonal line harbor HACs, matching this property of our prior generations of HACs (*11*, *12*, *14*, *16*). Four of the monoclonal lines (colored data points in graph in Fig. 2B) were subjected to three independent HAC stability assays over a month of growth in the absence of antibiotic selection (Fig. 2C). The average daily HAC loss rate of 0.011 +/-0.006 (Fig. 2C) was similarly low as those we and others have reported (*12*, *16*, *28*).

**Figure 2:**
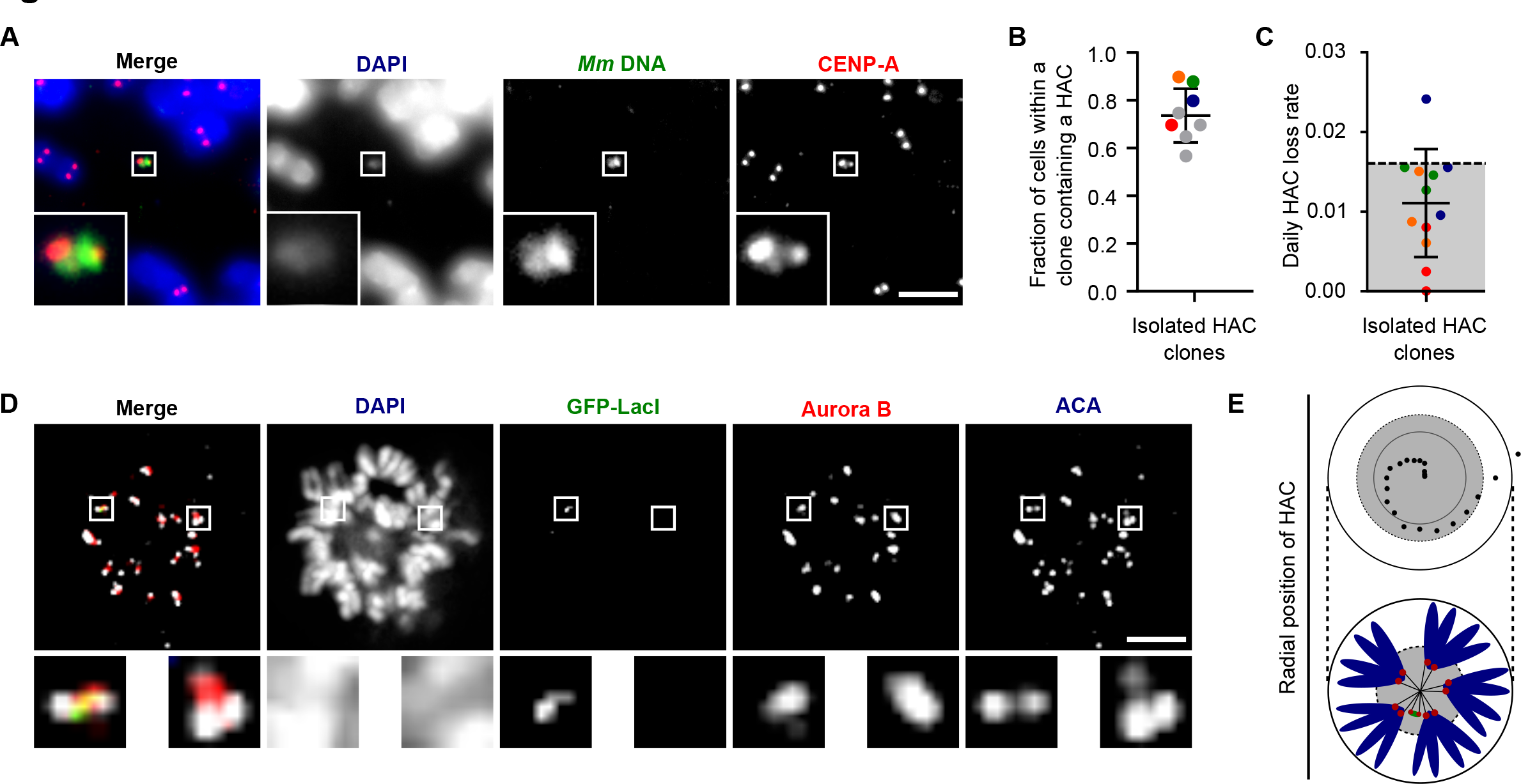
YAC-*Mm*-4q21^LacO^-based HACs are inherited as autonomous chromosomes with functional kinetochores and robust CPC recruitment. A) Representative image of a single copy HAC that has been isolated in a monoclonal cell line. Inset: 5x magnification. Bar, 5 μm. B) Quantification of fraction of spreads with a HAC in monoclonal cell lines. The mean (+/-SD) is shown. C) Quantification of HAC loss rate after culturing without selection for 30 days. The mean (+/-SD) is shown. Experiments are color coded to correspond to the clones shown in panel B. Grey shading indicates the range of loss rates for prior generations of HACs (*12*, *16*, *28*). D) Representative image of HACs synchronized in mitosis showing Aurora B and ACA. The image shows 8 0.2 μm z-projected stacks (see also Fig. S4 for centromere delineation in the z-dimension). Inset: 5x magnification. Bar, 5 μm. E) The radial position of HACs was measured relative to endogenous centromeres. The position of 20 HACs, each endogenous centromere and the center of DNA mass was measured. The distance between HAC or endogenous centromere and the center of DNA mass was calculated. The distance of each HAC from the center was normalized based on the total length across (i.e. the diameter) of mitotic chromosomes. The inner black circle represents the mean radial position of endogenous centromeres, while the dotted line represents one standard deviation from the mean. An illustration is shown below the graph.

To further interrogate our initial hypothesis that a larger initial HAC construct would confer full centromere function, we assessed the mitotic recruitment of a key component of the inner centromeric error correction mechanism, the Aurora B kinase (*18*). The region that harbors both the kinetochore-forming, CENP-A-containing chromatin and the inner centromere (including the conventional chromatin containing histone H3, but decorated with H3^T3phos^ and H2A^T121phos^ modifications for recruiting the Chromosome Passenger Complex [CPC] that includes Aurora B); (*29–31*) spans several Mbp on natural chromosomes (*32*). The inner centromere in metazoans is not a single thread of chromatin but rather thought to be a densely packed region that spans a linear distance of 500-1000 nm between sister centromeres (*33*, *34*). Given the high-fidelity of transmission of our HACs (Fig. 2C), we predicted that they are sufficiently large to generate a robust inner centromere that recruits Aurora B. In order to have the necessary dispersion of chromosomes in the mitotic cell for robust detection of the HAC via expression of GFP-LacI, we induced the formation of monopolar spindles and assessed both the kinetochore forming part of the paired sister HACs (with anti-centromere antibodies; ACA) and the inner centromere (with antibodies to Aurora B)(Fig. 2D,E, S4). In the vast majority of HACs, Aurora B was clearly detectable (detectable Aurora B was found in 33/35 HACs). Thus, our findings suggest that the increased size of YAC-*Mm*-4q21^LacO^ relative to prior HAC constructs permits a robust inner centromere without the need to undergo large-scale multimerization. This is critical since it endows YAC-*Mm*-4q21^LacO^ HACs with the ability to segregate in mitosis at high-fidelity alongside natural counterparts.

### Single-copy HACs

The small physical size of HACs formed from YAC-*Mm*-4q21^LacO^ (Fig. 1) raised the possibility that they can form without any multimerization at all. To test this notion, we sought a cytological approach that reports on copy number without the deformations that happen naturally when chromosomes are attached to and stretched by the spindle or otherwise confounded by mitotic chromosome condensation. Fortuitously, we found that nuclear envelope lysis during isolation of nuclei releases the small HACs formed with YAC-*Mm*-4q21^LacO^ that are subsequently efficiently separated from the rest of the genome via centrifugation (Fig. 3A-C). We harvested the top gradient fractions in the 10% sucrose layer (i.e. above the visible cell debris), and determined the location of CENP-A and the LacO sequences (Fig. 3D). We anticipated a single CENP-A focus on interphase HACs, even after replication, since sister centromeres are not separated into distinct foci on natural chromosomes until just prior to nuclear envelope breakdown near mitotic onset (*35*). LacO arrays, on the other hand, when present on repeated HAC constructs do not coalesce into a single focus (*16*). HACs were readily identified in these fractions, representing what to our knowledge is the first visualization of an individualized and functional metazoan chromosome in its decondensed, interphase form (Fig. 3D). In the vast majority of HACs, a single focus each of CENP-A and LacO sequences was present (Fig. 3D). We did not observe any HACs with a single CENP-A locus and more than one LacO locus. A small number of HACs harbored two CENP-A loci, consistent with them coming from cells were in late G2 or early mitosis (i.e. prior to sister chromatid separation) at the time of isolation. Importantly, each of these also had precisely two LacO foci (Fig. 3D). Unlike the single-copy YAC-*Mm*-4q21^LacO^ HACs, prior generations of HACs are large multimers that do not separate from endogenous chromosomes during nuclei isolation (*16*), but, in these HACs, CENP-A and LacO arrays are readily visualized on mitotic HACs in chromosome spreads (Fig. 3E). For the prior generation of HACs, the paired, replicated centromeres are visible as ‘double dots’ of CENP-A, whereas the LacO arrays are visible as numerous foci (Fig. 3E). Taken together with the earlier detection from uniformly small-sized HACs from populations of cells with nascent YAC-*Mm*-4q21^LacO^ HACs (Fig. 1B-D), our interphase HAC experiments (Fig. 3D,E) indicate that the new HACs are formed and maintained without multimerization.

**Figure 3:**
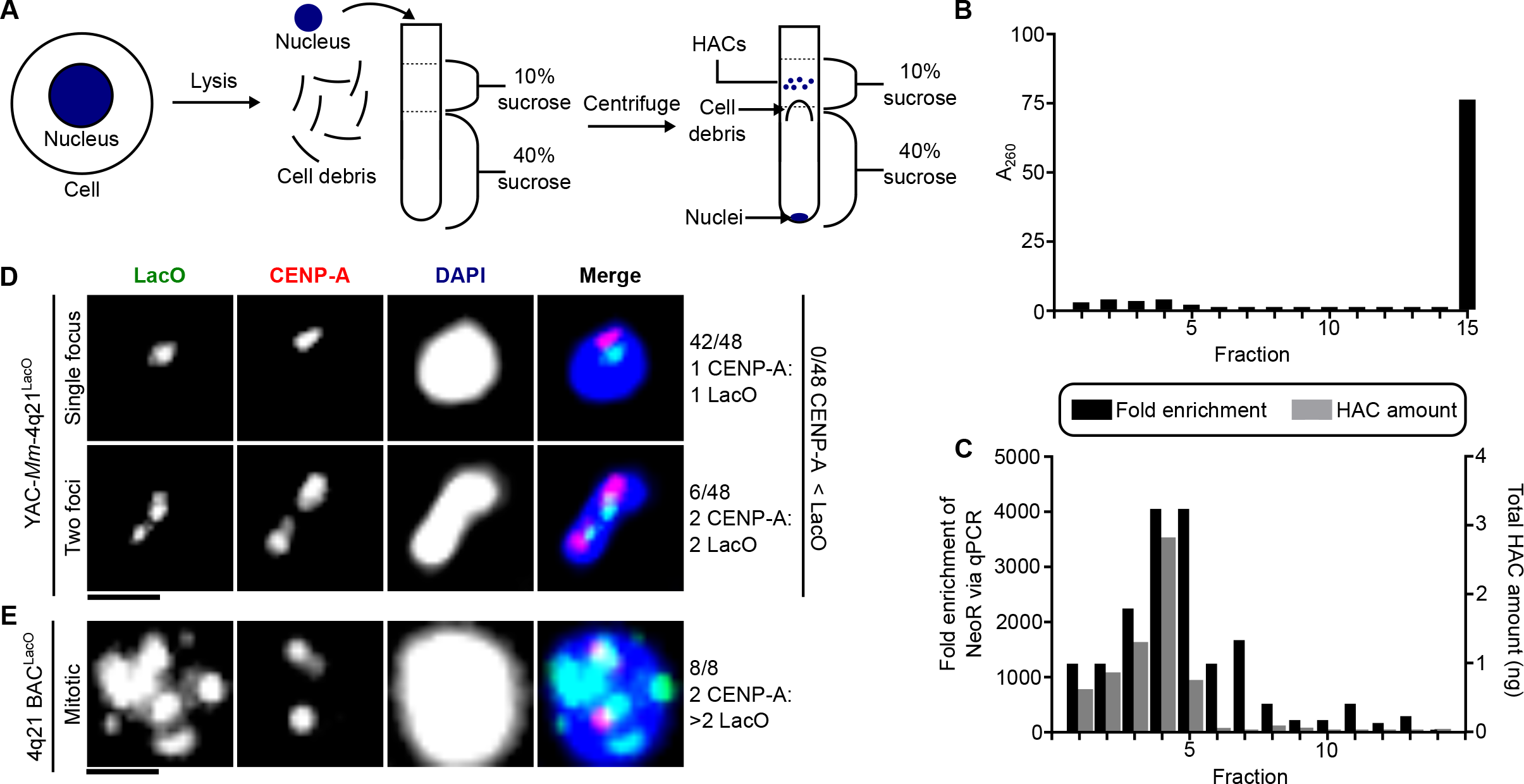
YAC-*Mm*-4q21^LacO^ HACs are functional as single copy DNA. A) Schematic of approach used to enrich single copy HACs. B) A_260_ measurements of fractions collected from a sucrose gradient from top (fraction 1) to bottom (fraction 14) as well as pelleted nuclei (fraction 15). Fraction 15 was diluted 33.3 x relative to other samples to acquire a reading in the measurable range (dilution corrected values are plotted). C) Enrichment of HAC DNA compared to endogenous chromosomal DNA. The HAC DNA concentration is also shown. D) Representative image of HACs isolated by sucrose gradient with either a single or two foci of LacO. The proportion of HACs with a single or two foci is noted. HACs with two LacO foci also had two CENP-A foci suggesting that they are mitotic. Bar, 1 μm. E) Representative image of a multimerized HAC (Clone 27 from (*16*)) from mitotic chromosome spreads. Bar, 1 μm.

We next assessed the size and topology of functional YAC-*Mm*-4q21^LacO^ based HACs (Fig. 4). This is important because earlier generations of HACs typically formed in a manner accompanied by large-scale DNA sequence multimerization and even acquisition of portions (>100 kb) of host cell chromosomal DNA (*16*, *36*, *37*). YAC-*Mm*-4q21^LacO^ contains a single FseI site (Fig. S1C), and we found that two isolated HAC-containing cell lines required FseI digestion to enter a pulse-field gel (Figs. 4A; S5). This is consistent with well-established topological trapping of circular chromosomes prior to digestion (*38*). The mobility of the linearized HAC suggests that it has not lost or gained large fragments of DNA (Figs. 4A; S5). We compared this to a circular, multimerized BAC-based HAC (*16*) that has one FseI site per repeating ∼200 kb monomer (Figs. 4A; S5). Along with our cytological data indicating the HACs are single copy (Fig. 3), their behavior on pulse-field gels (Figs. 4A; S5) support the notion that they function and are inherited through cell divisions with the same single-copy circular nature in which they were initially constructed in yeast.

**Figure 4:**
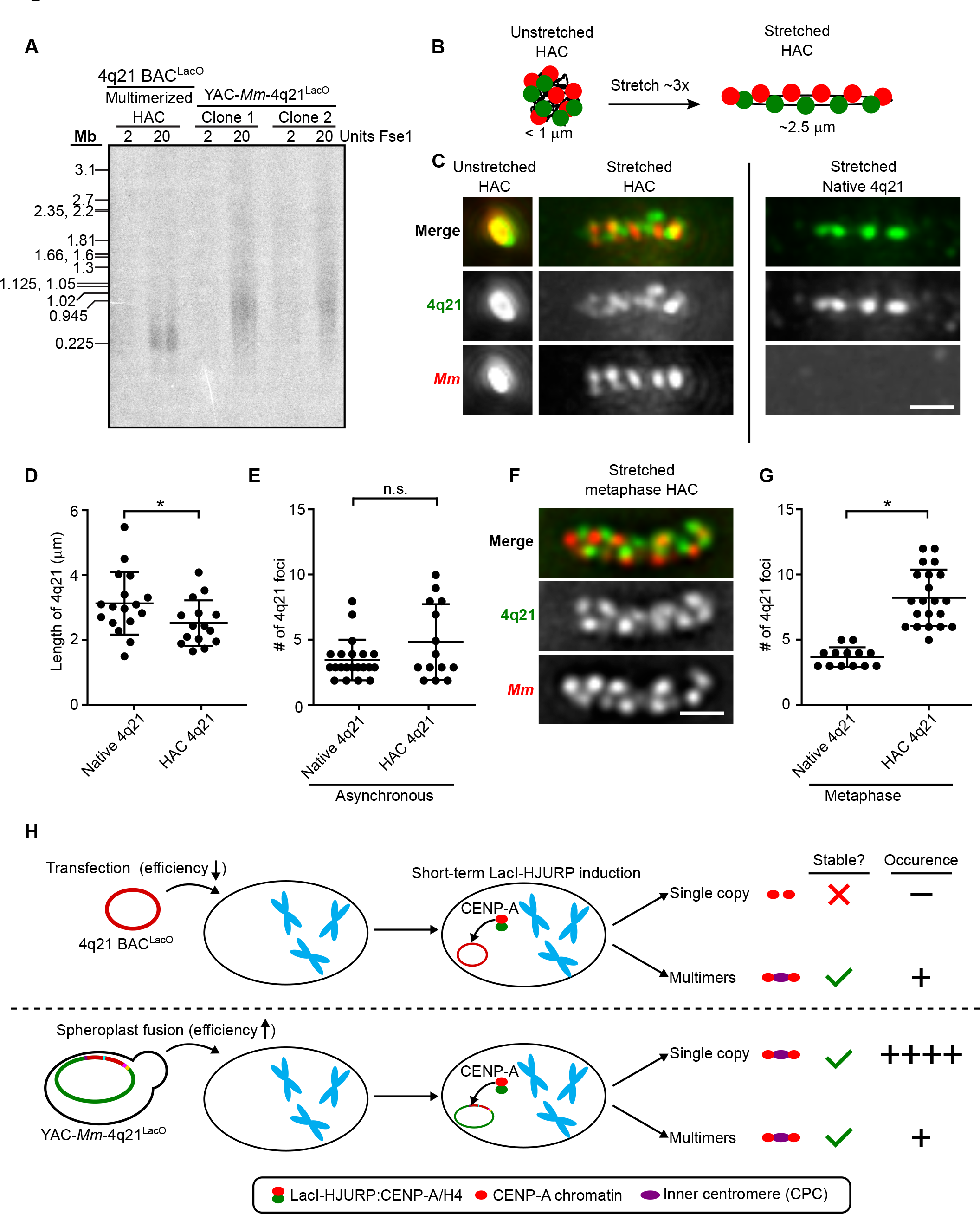
YAC-*Mm*-4q21^LacO^-based HACs are intact 760 kb circles with similar chromatin stretching properties as natural chromosomes. A) Southern bot analysis of the indicated HAC lines using a LacO probe. B) Schematic showing extent of stretching HACs in our experiments (panels C-G), with indicated regions detected by FISH. The number of foci shown is in the range predicted by prior stretching experiments with natural chromosomes, with actual outcomes measured in panels C-G and Fig. S6. C) Representative images of an unstretched and stretched HAC with both the 4q21 and *M. mycoides* sequence labeled via FISH compared to endogenous 4q21 in asynchronous cells. Bar, 1 μm. D) Quantification of the length of 4q21 FISH in the HAC and the endogenous chromosome after stretching chromatin in asynchronous cells. The mean (+/-SD) is shown. p < 0.05 based on an unpaired, two-tailed t-test. E) Quantification of the number of foci from 4q21 FISH in the HAC and 4q21 FISH in the endogenous chromosome after stretching chromatin in asynchronous cells. The mean (+/-SD) is shown. p value > 0.05 based on an unpaired, two-tailed t-test and is marked as not significant (n.s.). F) Representative images of a stretched HAC with both the 4q21 and *M. mycoides* sequence labeled via FISH after enriching for cells in metaphase. Bar, 1 μm. G) Quantification of the number of foci from 4q21 FISH in the HAC and the endogenous chromosome after stretching chromatin and enriching for cells in metaphase. The mean (+/-SD) is shown. p < 0.0001 based on an unpaired, two-tailed t-test. H) Model illustrating how construct size influences HAC formation outcomes.

To cytologically examine the shape and nature of the chromatin assembled on the HAC, we employed a well-established chromatin stretching approach (*33*, *39*) with which we could monitor the 182 kb region of 4q21 present on the HACs and at the endogenous locus on chromosome 4 in the same samples (Fig. 4B-G). The degree of stretching we achieve is about 3-fold, making the circle fold back upon itself (Figs. 4B,C; S6). Stretching at either locus maintains large blocks (roughly 40 kb) of chromatin linked by highly stretched regions with little or no detectable FISH signal (Figs. 4C and S6). Indeed, the overall length is decreased slightly in the HAC (3.13 +/-0.94 versus 2.52 +/-0.68 μm; Fig. 4D), while the number of foci produced by stretching of the native locus and HAC is similar (3.6 +/-1.5 versus 4.9 +/-2.8; Fig. 4E).

Interestingly, the number of 4q21 foci observed on the HAC varied greatly revealing the possible existence of two populations of HACs (Fig. 4E). We reasoned that the HAC would be visualized as a single chromatid early in the cell cycle (G1 and early S phase), whereas it would be visualized as paired chromatids later in the cell cycle (late S, G2, and M). On the other hand, the natural 4q21 locus would be visualized in stretching experiments as a single chromatid, even late in the cell cycle, since each natural chromosome arm location would make its own chromatin fiber. To test this notion, we enriched cells in mitosis, prior to sister chromatid separation, and found that the average number of 4q21 foci in the HAC increased (8.2+/-2.1) (Fig. 4 F,G) relative to those from asynchronous cell populations (Fig. 4E), whereas the endogenous 4q21 locus has a similar number of foci after stretching in both instances (Fig. 4E,G and S6). We note that the *M. mycoides* chromatin appears to be relatively resistant to stretching, since there is a similar number of foci and length (Figs. 4C-G and S6) despite its twice longer DNA sequence than 4q21 on YAC-*Mm*-4q21^LacO^. A likely explanation is that the AT-rich *M. mycoides* sequence has denser chromatin relative to the human 4q21 sequence. HAC stretching supports the notion of a chromatinized circular topology supporting propagation and inheritance in mitotically-dividing cells.

## Discussion

*De novo* HACs, like the ones we advance in this paper, are the only viable platform to generate a new mammalian chromosome where the entire DNA sequence can be designed in the lab. This presents an extremely wide horizon of possibilities for downstream biological and applied uses, and we report a system to create HACs that faithfully exist in their functional form as a single copy. The advances are centered on the portion of the chromosome, the centromere, that controls segregation of the HAC at cell division. Single-copy HAC formation requires establishment of a high local density of CENP-A nucleosomes that can self-propagate alongside natural centromeres (*16*), but this is not sufficient for centromere function. Rather, an entirely different chromatin domain, the inner centromere, must function in mitotic quality control and control of sister chromatid cohesion. The HACs formed from the YAC-*Mm*-4q21^LacO^ are large enough to harbor robust CENP-A arrays and inner centromeric chromatin (Figs. 2 and 4H).

The overhauled HAC cell delivery approach via spheroplast fusion was necessary because the initial construct is too large to reliably purify and transfect, and has the benefit of being far more efficient than HAC construct transfections. YACs from spheroplasts will be packaged into chromatin, which may also contribute to the high rate of HAC formation upon introduction into mammalian cells. The overall high efficiency of HAC delivery and formation upon moving to spheroplast fusion-based delivery has important practical implications for the development and testing of specific features of HACs. In prior generations of HACs, rigorous testing of a modest number of constructs or cell lines requires the isolation and subsequent cytological assessment of hundreds of cell lines that are cloned weeks after initial HAC construct transfection (*16*, *27*), since the initial selected cell populations are so sparse and HAC formation is so inefficient. With YAC-*Mm*-4q21^LacO^, we can measure high HAC formation efficiency in rapidly generated cell populations, without the requirement to generate any clonal lines. To measure the prior generation HACs with a similar level of repetition and statistical power as in one of our YAC-*Mm*-4q21^LacO^ experiments with four different conditions (Fig. 1D) would have required generating a minimum of 240 cell lines with prior generations of HACs. One can easily envision a multitude of HAC features –including their genetic cargo and recipient cell types—that are attractive to test in the future. The YAC-*Mm*-4q21^LacO^ approaches we report eliminate the initial need for isolating hundreds of cell lines at the outset for each new derivative.

The innovations in HACs described in this paper promise to bring artificial, synthetic chromosomes towards their potential in delivering useful cargos for biomedical and industrial applications. Since both the CENP-A-based epigenetic centromere specification mechanism and the inner centromeric dimensions and molecular constituency (e.g. sister centromere cohesion components and CPC) are common to diverse eukaryotic species, including in many agriculturally important plants, we envision that YAC-*Mm*-4q21^LacO^-based artificial chromosomes will be readily modified and extended into many useful biological systems. The YAC-*Mm*-4q21^LacO^ system also extends the capacity for HACs to advance our understanding of natural chromosomes. In other words, HACs can be used as testing grounds for designing what nature has evolved to ensure the stability of our genome between cell divisions and from one organismal generation to the next, as well as designing the functional features that govern chromosome “outputs” in gene expression programs and epigenetic regulation. For instance, the methodology we developed to isolate an intact HAC from nuclei (Fig. 3), notably extends HAC technology as attractive vehicles to visualize and probe principles of chromatin organization on individual interphase chromosomes. In summary, the advancements made in this study to HAC design, delivery, formation, and function will expedite both discovery-based and applied genome science.

## Methods

### YAC construction

A total of 6 fragments were prepared for TAR cloning to make a HAC forming YAC construct. Two linker fragments were ordered from IDT. The remaining two were PCR amplified from 4q21 BAC^LacO^ and a vector containing the URA3 gene and mCherry under a CMV promoter. The 4q21 BAC^LacO^ was restriction digested into two fragments with Mre1 and Nru1 prior to insertion into yeast.

Yeast (strain VL6-48N) cells containing a YAC with *M. mycoides* genomic DNA were transformed with a plasmid, pDB18-cas9-CRISPR, containing an expression cassette for guide RNAs to cut the yeast construct and Cas9. Yeast were grown in 30 ml SD-HIS overnight at 30 °C. Yeast cells were centrifuged at 1800 x g for 3 min. Cells were resuspended to an OD_600_ of 0.4 in YPG. Yeast cells were grown for 6 h to an OD_600_ of 1. Then, 1.5 ml of the culture was centrifuged at 14,000 rpm for 15 s with a microfuge. The supernatant was removed and resuspended in 1 ml 0.1 M LiOAc. Cells were centrifuged again at 14,000 rpm for 15 s before resuspending in 1 ml 0.1 M LiOAc. Cells were incubated at 30 °C for 30 min. Cells were centrifuged at 5,000 rpm for 3 min. Cells were resuspended in 50 µl 0.1 M LiOAc in 1x TE, and add 5 µl denaturated carrier DNA (10 ug/ml sheared salmon sperm DNA) and 10 µl DNA insert mix. With mixing between each addition, 500 μl of 40% PEG 4000 and 56 μl of DMSO were added to the solution. The solution was incubated for 30 min at 30 °C and then 25 min at 42 °C. The solution was spun at 5,000 rpm for 3 min before resuspending in 100 μl dH_2_O and plating on SD-HIS-URA plates.

### Junction PCR to assess YAC

Genomic DNA was prepared as described (*40*) by first growing yeast overnight in a 5 ml culture to and OD_600_ > 0.4. Then, 200 μl of yeast was centrifuged at 2000 x g for 3 min at 4°C. Yeast was resuspended in 100 ul 200 mM LiOAc with 1% SDS and incubated for 5 min at 70°C. Yeast DNA was first extracted with Phenol-Chloroform-Isoamyl alcohol before precipitating with isopropanol and resuspending in elution buffer (EB; Qiagen). YAC containing yeast were assessed by PCR using Q5® Hot Start High-Fidelity 2X Master Mix (M0494S). The primers sets used were: (1: Fwd: 5’-gtaccaccgcaactttcttg-3’, Rev: 5’-cggcgcagtttctgagaag-3’, 2: Fwd: 5’-TATTGGTGAACCAGTGGG-3’, Rev: 5’-CCTTGTTCAACACGTAATACTG-3’, 3: Fwd: 5’-CCGTAATATCCAGCTGAACG-3’, Rev: 5’-CAGCCAAGATATCAGCATCA-3’)

### Sequencing to assess the YAC

For short read sequencing 3 μg yeast genomic DNA was digest with NEBNext dsDNA Fragmentase (NEB; # M0348S) in Fragmentase Reaction Buffer v2 at 30 °C for 30 min before stopping the reaction with EDTA. Sequencing libraries were generated and barcoded for multiplexing according to Illumina recommendations with minor modifications. Briefly, 10 ng DNA was end-repaired and A-tailed. Illumina TruSeq adaptors were ligated, libraries were size-selected to exclude polynucleosomes, and the libraries were PCR-amplified using KAPA DNA polymerase. All steps in library preparation were carried out using New England BioLabs enzymes. Resulting libraries were submitted for 75-bp, single-end Illumina sequencing on a NextSeq 500 instrument.

For Oxford Nanopore Technologies (ONT) long-read sequencing of yeast harboring YAC-*Mm*-4q21^LacO^, genomic DNA was first incubated in 25 μg/ml RNase for one h at 37 °C and then extracted with Phenol-Chloroform-Isoamyl alcohol before precipitating with isopropanol. Once the DNA was fully resuspended in EB the following day, DNA was sheared with a g-tube (Covaris) we prepared the DNA for ONT long-read sequencing using the ONT ligation sequencing kit (ONT; # SQK-LSK112), following the manufacturer’s instructions. The library was loaded onto a primed FLO-MIN106 R9.4.1 flow cell for sequencing on the MinION. All ONT data was basecalled with Guppy 3.6.0 with the HAC model.

To generate the sequence of the YAC, our expected input sequence, short (because of the high read accuracy) and long (to identify any major sequence rearrangements) read sequencing data was input into the EPI2ME pipeline wf-bacterial-genomes v0.2.12. The draft assembly output was used as a template for two kinds of manual revisions. First, in several places long reads spanned a gap with no coverage, the regions with no coverage were deleted to allow continuous coverage of those long reads. Second, in two locations, there were insertions not represented by the existing sequence map. Thus, reads with alignment to those regions were identified and the sequence from those reads not on the existing map was added to the assembly. Alignments were performed via the EPI2ME pipeline wf-alignment v0.3.3.

### Spheroplast fusions

Yeast harboring YAC-*Mm*-4q21^LacO^ were grow overnight to saturation in a 5 ml culture of SD-URA. This culture was diluted to 50 ml in SD-URA and grown for 7-8 h at 30 °C to an OD_600_ of 0.8-1.0. Yeast were centrifuged at 3,000 rpm for 3 min in an A-4-62 swing bucket rotor (Eppendorf) and resuspended in 20 ml 1 M sorbitol and incubated at 4 °C overnight (<18 h). Yeast cells were spun down at 3,000 rpm for 3 min and resuspended in 20 ml SPEM (1 M sorbitol, 10 mM EDTA, 10 mM sodium phosphate at pH 7.4). 40 μl of BME and 60 μl of zymolase (stock solution of 200 mg Zymolase 20-T resuspended in 9 ml H_2_O, 1 ml 1 M Tris pH 7.5, 10 ml 50% glycerol and stored at −20 °C) were added to the yeast solution. Yeast were incubated for 1 h at 37 °C to digest the cell wall. The success of spheroplasting was assessed by measuring the OD^600^ of solution diluted 1:10 in 1 M sorbitol and 1:10 in 2% SDS. The 37 °C incubation was continued until the OD^600^ ratio was >10. 30 ml of 1 M sorbitol at 4 °C was added before spinning at 1,800 rpm for 8 min at 4 °C. The pellet was resuspended in 20 ml 1 M sorbitol before adding an additional 30 ml of 1 M sorbitol. The solution was centrifuged at 1,800 rpm for 8 min at 4 °C. The pellet was resuspended in 1 ml of STC (1 M sorbitol, 10 mM CaCl_2_, 10 mM Tris pH 7.5) and incubated at RT for 10-60 min.

Tissue culture cells were processed in parallel to yeast. The day of fusion, 10 μl of 50 mg/ml S-trityl-L-cysteine (STLC) was added to 70-80% confluent plate of HT1080 or U2OS cells for 6 h. Cells were trypsinized and neutralized with DMEM with 4.5 g/L D-Glucose and L-Glutamine (Gibco) before counting on a hemacytometer and spinning at 1500 rpm for 5 min at RT. Cells were resuspended in PBS to a concentration of 6 x 10^5^ cells/ml. The concentration of yeast cells was determined by assuming 2 x 10^7^ cells per OD per ml. 3 x 10^5^ mammalian cells and 9 x 10^7^ yeast cells were mixed in an Eppendorf tube and incubated for 5 min at RT. The mixture was spun at 4,000 rpm for 30 s on a tabletop centrifuge. The pellet was resuspended in 45% PEG, 10% DMSO in 75 mM HEPES at pH 8.0 and incubated for 5 min at RT. The reaction was quenched by adding 1 ml of DMEM to solution before spinning at 4000 rpm for 30 s. The mixture was resuspended in 1 ml of DMEM before adding the mixture to a 6-well plate containing 2 ml of DMEM with 4.5 g/L D-Glucose and L-Glutamine (Gibco) supplemented with supplemented 10% FBS, 100 U/mL penicillin, and 100 μg/ml streptomycin. Cell lines were maintained at 37 °C in a humidified incubator with 5% CO_2_ (all HT1080 and U2OS cells were cultured in these conditions unless otherwise stated). After 3-4 h and cells have adhered to the plate, the media was replaced with fresh media containing 2 μg/ml doxycycline. The media was replaced again the following morning.

After fusion, cells were incubated in DMEM with 2 μg/ml dox for 48 h. Next, cells were trypsinized and moved to a 10 cm plate and cultured in DMEM supplemented 10% FBS, 100 U/mL penicillin, and 100 μg/mL streptomycin with 333 μg/ml G418-S for 8 days. Cells were then moved to a lower concentration of G418-S (150 μg/ml) and, after three days, were processed further for IF-FISH, frozen down, or isolated into single clones.

### Isolating monoclonal cell lines harboring HACs

Polyclonal HAC lines were first trypsinized before quenching with DMEM. Cells were centrifuged at 1,500 rpm for 5 min before being washed once with PBS. Cells were counted using a hemacytometer and centrifuged at 1,000 rpm for 3 min before resuspending in 10 ml PBS with 1 mM EDTA. The cells were centrifuged and resuspended once more to wash them. Cells were centrifuged once again at 1,000 rpm for 3 min before resuspending in PBS supplemented with 1 mM EDTA and 1% BSA to a final concentration of 10^6^ cells/ml before being transferred to a 5 ml sterile polystyrene tube. Single cells were sorted in wells of a 96-well plate using a FacsJAZZ sorter. Cells were cultured for ∼2-3 weeks in 50% DMEM and 50% conditioned media. Conditioned media was made by culturing HT1080 or U2OS cells overnight, collecting media and filtering with a 0.22 μm filter. After colonies were visible, cells were scaled up to a 24-well, 6-well and then 10 cm plate while culturing in DMEM (need to indicated with what kind of serum/concentration) supplemented with 150 μg/ml G418-S. Clones were assessed for the presence of HACs via IF-FISH on metaphase spreads.

### IF-FISH on metaphase spreads

IF-FISH was performed as described (*41*) with some modifications. HT1080 cells were treated with 50 μM STLC for 2-4 h to arrest cells during mitosis. Mitotic cells were blown off using a transfer pipette and swollen in a hypotonic buffer consisting of a 1:1:1 ratio of 75 mM KCl, 0.8% NaCitrate, and 3 mM CaCl_2_, and 1.5 mM MgCl_2_ for 15 min. 3 x 10^4^ cells were cytospun at 1500 rpm on high acceleration in a Shandon Cytospin 4 onto an ethanol-washed positively charged glass slide and allowed to adhere for 1 min before permeabilizing with KCM buffer for 15 min. Cells were blocked for 20 min in IF block buffer (2% FBS, 2% BSA, 0.1% Tween-20, and 0.02% NaN_3_) before incubating for 45 min at RT with a monoclonal anti-CENP-A antibody (Enzo; ADI-KAM-CC006-E) diluted 1:1000 in IF block buffer. Slides were washed 3 x 5 min in KCM buffer. Slides were incubated for 25 min at RT with Cy3 conjugated to donkey anti-mouse diluted 1:200. Slides were washed 3x in KCM for 5 min at RT. Slides were fixed in 4% formaldehyde in PBS, before washing 3x in dH_2_O for 1 min each. Slides were incubated with 5 μg/ml RNAseA in 2x SSC at 37 °C for 5 min. Cells were subjected to an ethanol series to dehydrate the cells and then denatured in 70% formamide/2x SSC at 77C for 2.5 min. Cells were dehydrated again with an ethanol series.

Biotinylated DNA probe was generated using purified *M. mycoides* DNA with a Nick Translation Kit (Roche; 10976776001) according to the manufacturer’s instructions, purified with a G-50 spin column (Illustra), and ethanol-precipitated with salmon sperm DNA and Cot-1 DNA. Precipitated BAC^LacO^ DNA or LacO plasmid was suspended in 50% formamide/10% dextran sulfate in 2x SSC and denatured at 77 °C for 5-10 min before being placed at 37 °C for at least 20 min. 300 ng DNA probe was incubated with the cells on a glass slide at 37 °C overnight in a dark, humidified chamber. The next day, slides were washed 2x with 50% formamide in 2x SSC for 5 min at 37 °C (45 °C for repetitive LacO FISH probe). Next, slides were washed 2x with 2x SSC for 5 min at 37 °C (45 °C and 0.1x SSC for repetitive lacO FISH probe). Slides were blocked with 2.5% milk in 4x SSC with 0.1% Tween-20 for 10 min. Cells were incubated with NeutrAvidin-FITC (ThermoFisher Scientific; 31006) diluted to 25 μg/mL in with 2.5% milk in 4x SSC with 0.1% Tween-20 for 10 min for 1 h at 37 °C in a dark, humidified chamber. Cells were washed 3x with 4x SSC and 0.1% Tween-20 at 45 °C, DAPI-stained, and mounted on a glass coverslip with Vectashield (Vector Labs). Slides were imaged on an inverted fluorescence microscope (Leica DMI6000B) equipped with a charge-coupled device camera (Hamamatsu Photonics ORCA AG) and a 100x 1.4 NA objective lens.

A “small HAC” designation was given if the cell contained a chromosome in which the FISH signal colocalized with CENP-A signal, was not overlapping an endogenous centromere, and the maximum diameter was < 1.0 μm. A “large HAC” designation was given if the cell contained a chromosome in which the FISH signal colocalized with CENP-A signal, was not overlapping an endogenous centromere, and the maximum diameter was > 1.0 μm. An ‘‘integration’’ designation was given if the cell contained a chromosome in which FISH probe signal localized to the DAPI-stainable region on the chromosome but did not colocalize with CENP-A signal; and a ‘‘no signal’’ designation was given if the cell did not contain a BAC probe signal on any DAPI-stainable region or colocalized with CENP-A signal.

For polyclonal HAC lines, 50 spreads were counted for each experimental condition and each HAC assay was performed in triplicate. The fraction of HACs with each designation was determined by dividing by 50. For isolated clones, 20 spreads were imaged and a clone was considered a HAC line if >20% of spreads contained a “small HAC” and no integrations or large HACs were present. The fraction of HACs in the isolated clone was determined by dividing the total number of “small HACs” by 20.

### Lentivirus production

HA-LacI or EGFP-LacI lentivirus was produced by co-transfecting the HA-LacI or EGFP-LacI lentiviral plasmid and two packaging plasmids, pMD2.G and psPax2 (Addgene plasmids #12259 and #12260, respectively), into 293GP cells (*42*) and harvesting the media 48 h later. Specifically, a 10 cm plate of 50%–80% confluent 293GP cells was transfected with 6 μg of DNA (3 μg of the HA-LacI lentiviral vector, 750 ng pMD2.G, and 2.25 μg psPax2) and 18 μL of FuGENE 6 (Promega). The culture medium was changed 6-24 h later. 48 h post-transfection, the culture medium was harvested, filtered through a 0.45 μm filter, and stored at −80 °C.

### IF of mitotic cells

HAC-containing cells were plated in a 6-cm plate (in the presence of 150 μg/ml G418-S) and allowed to adhere to the bottom of the plate. The next day (when cells were 20%–30% confluent), the culture medium was replaced with fresh medium containing 500 ml of eGFP-LacI lentiviral supernatant and 18 μg polybrene (Specialty Media, TR-1003-G). 24 h later, the culture medium was changed to remove the lentiviral particles and polybrene. 48 h after transduction cells were seeded on an 18 x 18 mm^2^ polystyrene coated coverslip. The following day, coverslips were transferred to a 6-well plate containing preheated PBS. PBS was removed via aspiration and cells were fixed in 1 ml of 4% formaldehyde in PIPES 60 mM PIPES, 25 mM HEPES, 10 mM EGTA, and 4 mM MgSO_4_·7H_2_0 at pH 6.9 with 1% Triton-X-100 was added to each well and incubated for 20 min at 37 °C. Formaldehyde solution was removed and fixing quenched with 2 ml 100 mM Tris pH 7.5 for 5 min. Slides were washed 3x in 2 ml PBS + 0.1% Tween for 5 min.

Slides were placed in IF Block (2% FBS, 2% BSA, 0.1% Tween-20, and 0.02% NaN_3_) for 20 min at RT. Slides were then incubated for 45 min at RT in a human ACA (Antibodies Inc.; 15-235) that we prepared by affinity purifying with recombinant CENP-A/H4 heterotetramers (*43*) and used at 0.74 μg/ml, mouse Aim-1 antibody (BD Transduction Laboratories; 611082) diluted 1:1,000 (serum), and rabbit anti-GFP antibody (made in-house) (*44*) used at 0.1 μg/ml in IF Block. Slides were washed 3x in 2 ml PBS supplemented with 0.1% Tween-20 for 5 min. Slides were then incubated for 25 min at RT in IF Block with Cy5 conjugated to donkey anti-human diluted 1:200, Cy3 conjugated to goat anti-mouse, and FITC conjugated to anti-rabbit. Slides were washed in 2 ml PBS + 0.1% Tween for 5 min before incubating in DAPI diluted 1:10,000 in PBS + 0.1% Tween-20 for 10 min. Coverslips were washed in PBS + 0.1% Tween-20, PBS, and then dH_2_O before mounting coverslips on slides with vectashield. Slides were imaged on an inverted fluorescence microscope (Leica DMI6000B) equipped with a charge-coupled device camera (Hamamatsu Photonics ORCA AG) and a 40x 1.4 NA objective lens. HACs were identified via the presence of GFP signal. Each HAC was determined to be Aurora B positive if Aurora B signal was at least 50% above background. The total fraction of Aurora B positive HACs was measured across three independent experiments.

### HAC retention assay

Four isolated HAC clones were cultured in the absence of G418-S selection for 60 days in triplicate. IF-FISH was performed at Day 0 and Day 30, and at least 20 cells were assessed for the presence of a HAC in each cell line at both time points. A daily HAC loss rate was determined using the following equation: N_30_ = N_0_ (1-R)^30^, where R is the daily HAC loss rate and N_0_ and N_30_ are the number of metaphase chromosome spreads containing a HAC at Day 0 and Day 30, respectively (*12*, *28*).

### Enriching HACs via a sucrose gradient

8 15-cm plates of cells harboring HACs were cultured to a confluence of 80-95%. Cells were centrifuged at 1,500 rpm for 5 min at 4 °C and resuspended in 30 ml of PBS. Cells were counted using a hemacytometer. Cells were centrifuged at 1,500 rpm at 4 °C. Keeping cells on ice, the cell pellet was resuspended in 0.32 M sucrose in 60 mM KCl, 15 mM NaCl, 5 mM MgCl_2_, 0.1 mM EGTA, 0.5 mM DTT, 0.1 mM PMSF, 1 mM leuptatin/pepstatin, 1 mM aprotinin, and 15 mM Tris pH 7.5. 2 ml of 0.32 M sucrose in 60 mM KCl, 15 mM NaCl, 5 mM MgCl_2_, .1 mM EGTA, 0.5 mM DTT, 0.1 mM PMSF, 1 mM leupeptin/pepstatin, 1 mM aprotinin, and 15 mM Tris pH 7.5 with 0.1% IGEPAL were added to 2 ml of cells and incubated for 10 min. The mixture was added onto a Sarsdedt tube containing 8 ml of 1.2 M sucrose in 60 mM KCl, 15 mM NaCl, 5 mM MgCl_2_, 0.1 mM EGTA, 0.5 mM DTT, 0.1 mM PMSF, 1 mM leuptatin/pepstatin, 1 mM aprotinin, and 15 mM Tris pH 7.5. The mixture was added slowly to avoid mixing the two layers of differing sucrose concentration. The sucrose gradient was centrifuged at 10,000 g, 20 min, 4°C at acceleration setting 9 and deceleration setting 5 with an SS-34 rotor in a Sorvall centrifuge. Individual 1 ml fractions were collected for qPCR analysis and the top ∼2.5 ml of solution (with care to avoid collecting cell debris) were collected for IF-FISH experiments.

### qPCR of sucrose gradient enriched HACs

DNA collected from sucrose gradient was first extracted with Phenol-Chloroform-Isoamyl alcohol before precipitating with isopropanol. DNA concentrations were determined via a Nanodrop. qPCR was performed in triplicate with 10 ng of initial DNA used in each reaction. qPCR amplification was detected using a 2x SYBR green master mix. Two primer sets were used, one with amplification of the CENP-A gene (present on endogenous chromosome) and another with amplification of the NeoR gene (present on the HAC). Nucleic acid amount was determined by an A^280^ measurement via a NanoDrop 2000 spectrophotometer. HAC enrichment was determined by the following calculation: fold enrichment = 1.81^([Ct sucrose fraction CENP-A gene – Ct sucrose fraction NeoR gene] – [Ct genomic DNA CENP-A gene – Ct genomic DNA NeoR gene]. HAC content in each fraction was determined by the following calculation: HAC DNA = 1.81^[Ct sucrose fraction NeoR-Ct genomic DNA NeoR]*[total DNA] * 760/6,270,000. Note that this assumes 1 HAC per cell and a diploid genome.

### IF-FISH on HACs isolated via sucrose gradient

HACS collected from the top of a sucrose gradient were cytospun onto slides and IF-FISH was performed as described above, but with the following modifications. Before cytospinning, 50 μl HAC solution was diluted in 450 μl H_2_O and incubated for 15 min. Next, during IF, slides were incubated at RT with CENP-A antibody for 2.5 h. HACs were identified on the slide via colocalization of DAPI (with size of the HAC DNA <2.5 μm and >.5 μm), CENP-A IF signal, and LacO FISH signal. The number of foci containing CENP-A signals and LacO signals were counted from two separate experiments.

### Southern blots

Genomic DNA from the indicated cell lines was prepared in agarose plugs by resuspending 5 x 10^6^ cells/ml in 0.8% agarose and digested overnight with Fse1 (NEB; R0588L) at 37 °C. Digested DNA was separated via CHEF electrophoresis (Bio-Rad, CHEF DR II System) at 3 V/Cm, 250 to 900 s, for 50 h. The blot was transferred to a membrane (Amersham Hybond-N+) and blot-hybridized with a 100 bp probe that binds to the LacO sequence (5’-TTGTTATCCGCTCACAATTCCACATGTGGCCACAAATTGTTATCCGCTCACAATTCCACATGTGGCCACAAATTGTTATCCGCTCACAATTCCACATGTG-3’). The LacO-specific probe was end labeled with ^32^P-γ-ATP for 1 h at 37 °C before cleaning with illustra ProbeQuant G-50 micro column (GE Healthcare; 28-9034-08). The blot was incubated for 2 h at 42 °C in hybridization buffer (ULTRAhyb™ Ultrasensitive Hybridization Buffer [Invitrogen; AM8669]). The probe was added to hybridization buffer and hybridized to the blot overnight at 38 °C. The blot was washed twice with 2x SSC with 0.5% SDS for 30 min at 42 °C. Finally, the blot was exposed to a phosphorimager screen for 2 weeks before imaging with an Amersham Typhoon.

### FISH on stretched chromatin fibers

Extended chromatin fibers were prepared and FISH was performed as described (*39*) with some modifications. The modified steps include the following: 5 × 10^4^ of HAC-containing cells were pelleted by centrifugation at 1000 g for 5 min at RT. The cell pellet was resuspended in 500 μl of hypotonic buffer (75 mM KCl) and incubated for 10 min at RT. Slides were then cytospun for 4 min at 800 rpm on high acceleration in a Shandon Cytospin 4 onto a poly-lysine coated glass slide. Slides were transferred quickly into a falcon tube containing freshly prepared salt-detergent lysis (SDL) buffer composed of 25 mM Tris-HCl (pH 9.5), 500 mM NaCl, 1 mM PMSF, and 1% Triton X-100. After 20 min of incubation at RT, slides were washed for 15 min in PBS supplemented with 0.1% Triton-X-100 and again in SDL buffer for 15 min before fixation with 3.7% formaldehyde. For experiments with mitotic enrichment for chromatin fiber stretching, colcemid was added to cell cultures at a final concentration of 0.1 μg/ml and incubated at 37 ^0^C in the presence of 5% CO_2_ for 3-4 h. Growth flasks were then gently tapped with the palm of the hand, dislodging mitotic cells from the surface. Mitotic cells were harvested and transferred into a 15 ml falcon tube and centrifuged at 1500 rpm for 5 min at RT. The pellet was resuspended in 0.5 ml PBS and cells were counted. An aliquot of cell suspension of concentration 1 x 10^5^ cells/ml was centrifuged at 1500 rpm for 5 min at RT. The pellet was resuspended in 1 ml of hypotonic buffer and incubated at RT for 15 min. An aliquot of 500 μl of cell suspension was loaded into cytospin funnel with poly-lysine coated slide and centrifuged at 1500 rpm for 5 min with cytospin set to high acceleration. One slide was carried through the fiber preparation protocol and the other slide proceeded through the mitotic spread protocol described above as a control slide to confirm successful mitotic chromosome enrichment.

## Acknowledgements

We thank our UPenn colleagues G. Birchak and M. Lampson for comments on the manuscript, and M. Gerace for assistance with preparing reagents.

## Author Contributions

C.W.G., D.M.B., J.I.G., and B.E.B. conceived the project. C.W.G., E.M., D.M.B., G.Y., and J.N.G. performed experiments. C.W.G., E.M., G.Y., J.N.G., P.H., and B.E.B. analyzed data. C.W.G., D.M.B., and G.A.L. generated reagents. C.W.G. and B.E.B. wrote the paper. All authors edited the paper. B.E.B. supervised the project.

## Data and Materials Availability

All sequencing data will be made publicly available at the time of publication on SRA. All data has been uploaded to SRA (PRJNA985068). All other data needed to evaluate the conclusions in this paper are present in the paper and/or the Supplementary Materials. The material used in this study are available from commercial sources of from the corresponding authors on reasonable request upon publication of the study.

---

**Figure S1:**
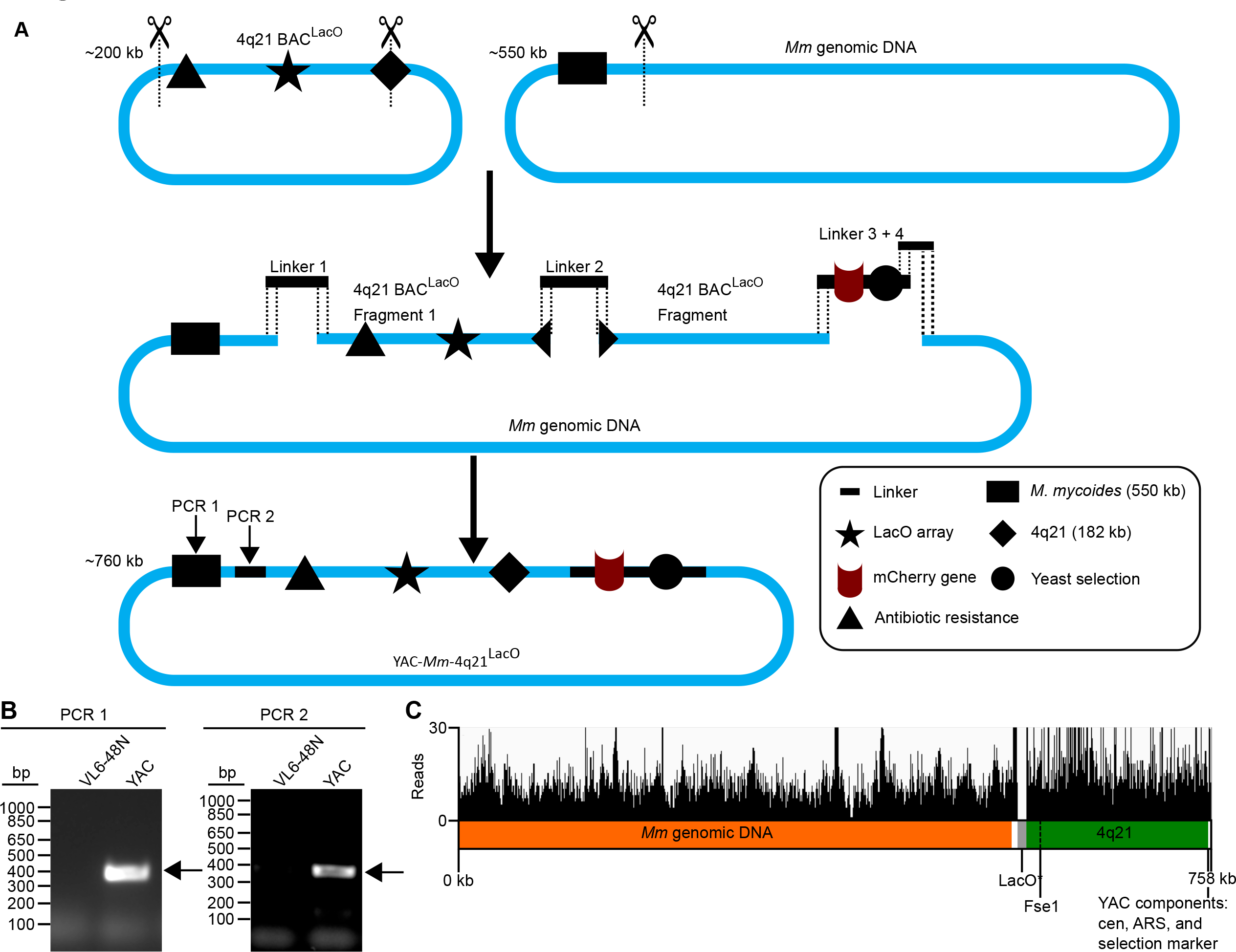
A 760 kb YAC construct with the necessary components for HAC formation was generated via TAR cloning. A) Schematic of YAC construct, *Mm*-4q21^LacO^, that was formed to generate single copy HACs. B) Junction PCR used to validate the YAC construct. C) Draft assembly (see Methods) of YAC-*Mm*-4q21^LacO^. Sequencing reads aligned to the YAC construct confirm presence of all components of the YAC except for the lacO array. A separate alignment of reads to the LacO array while allowing for multiple alignments confirmed the presence of the LacO array.

**Figure S2:**
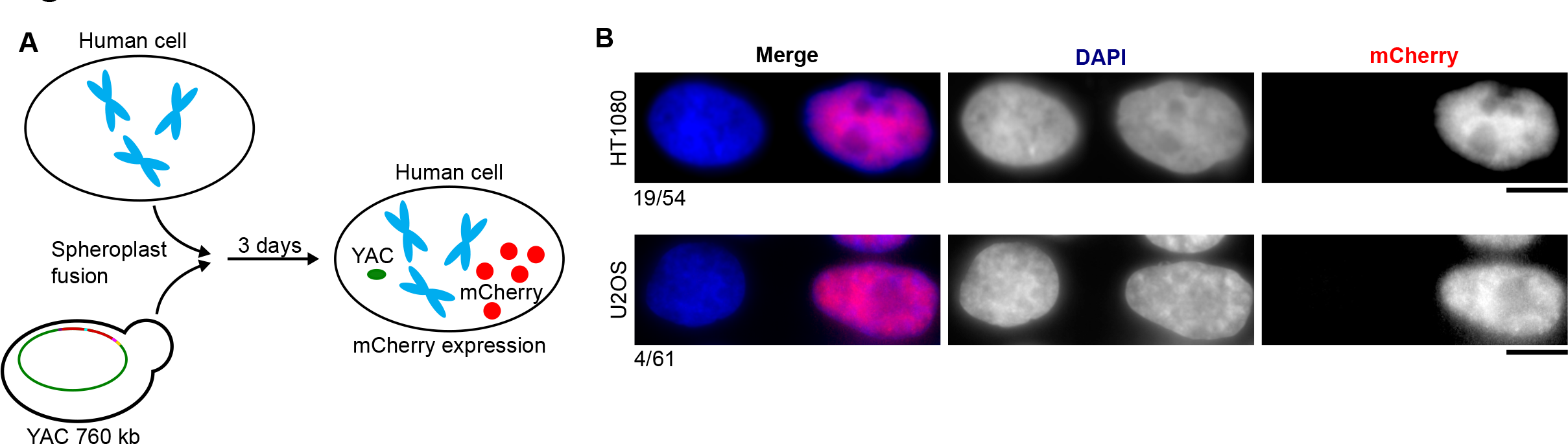
Yeast fusion approach is effective for delivering large DNA constructs in U2OS and HT1080. A) Schematic of approach to test for successful yeast fusion. B) Examples of successful yeast fusion with mCherry expression as well as the proportion of cells showing mCherry expression. The proportion of cells that were mCherry positive for each cell line is noted. Bar, 10 μm.

**Figure S3:**
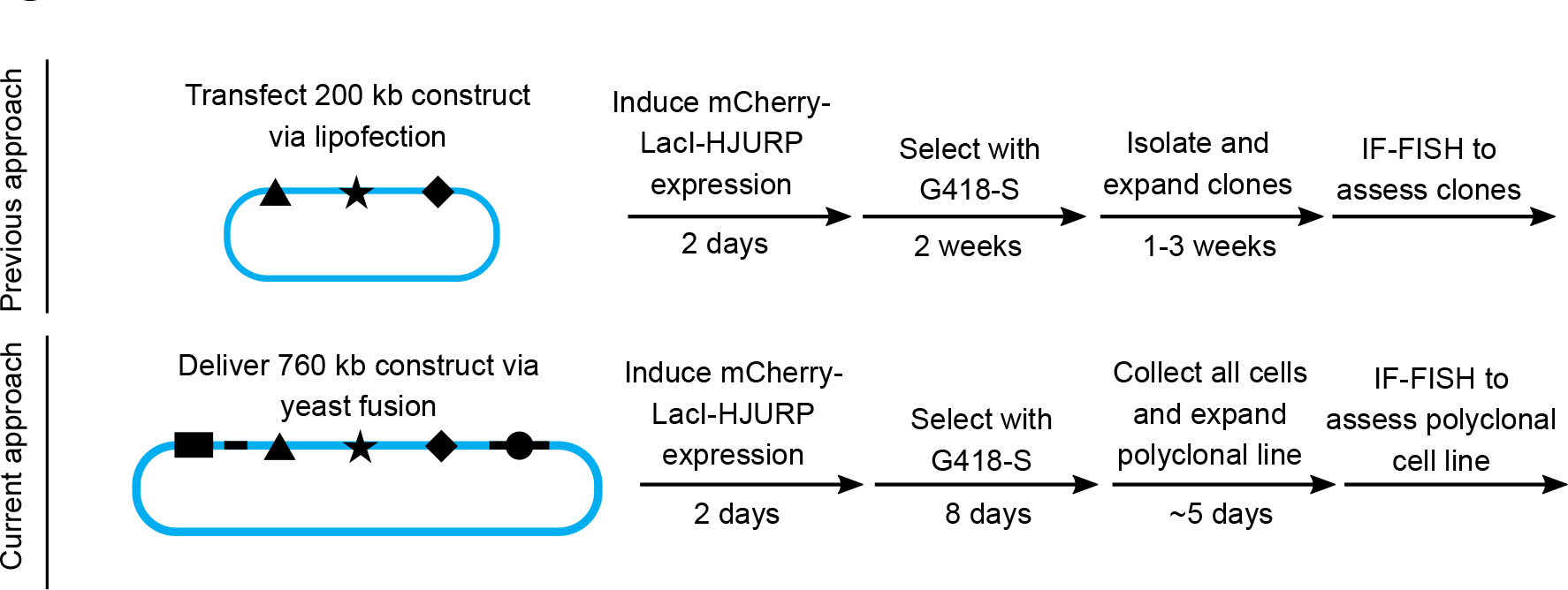
Schematic comparing the approach to generate HACs in prior generations (*16*) and the current approach.

**Figure S4:**
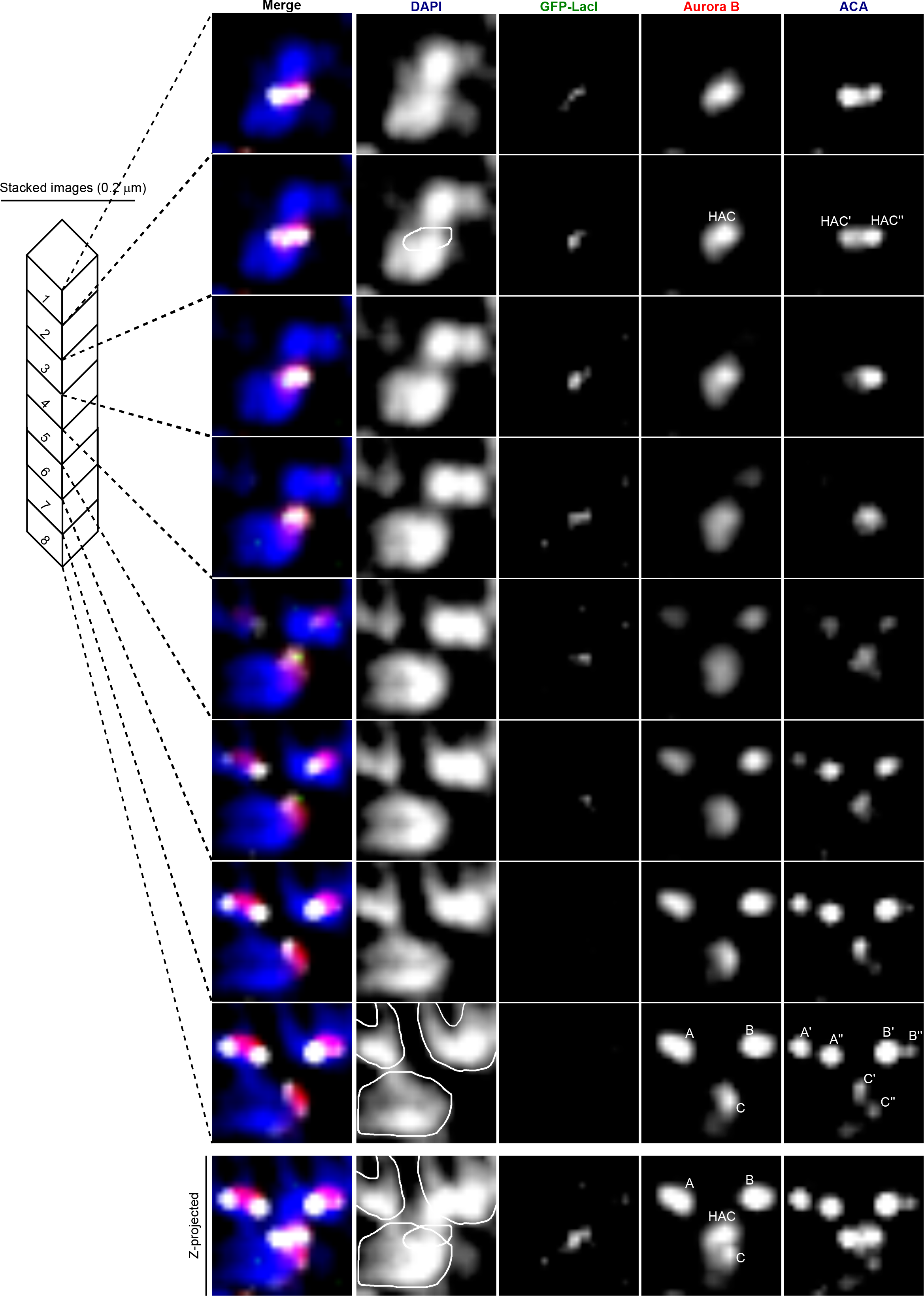
3-dimensional localization of HACs in monopolar mitotic cells. Mitotic chromosomes commonly overlap upon z-dimensional projections but are separable upon analysis of individual z-stacked images. This is especially important since DAPI staining is so heavily dominated by the natural chromosomes that are ∼100-fold larger than single-copy HACs. Three native chromosomes (labeled A, B and C) are immediately adjacent or overlapping to the HAC in the x and y dimensions but are ∼1 μm from the HAC in the z dimension. The close proximity of these chromosomes can account for the DAPI staining seen near to the HAC. In the z-stack of maximal Aurora B intensity, Aurora B from the HAC or natural chromosomes is labeled HAC, A, B or C and centromere double dots are labeled with HAC’, HAC’’, A’, A’’ B’, B’’, C’ or C’’. Additionally, the outlines of these chromosomes based on DAPI staining are shown. A z-projected image of all 8 stacks and all centromeres labeled is shown below the individual z-stacks. The peak Aurora B fluorescence is found in the following z-stack images: HAC, 2; A, 8; B, 8; C, 7. The peak of the two ACA foci is found for each centromere are found in the following z-stack images: HAC’, 1; HAC’’, 3; A’, 8, A’’, 8; B’, 8, B’’, >8; C’, 7; C’’, 8.

**Figure S5:**
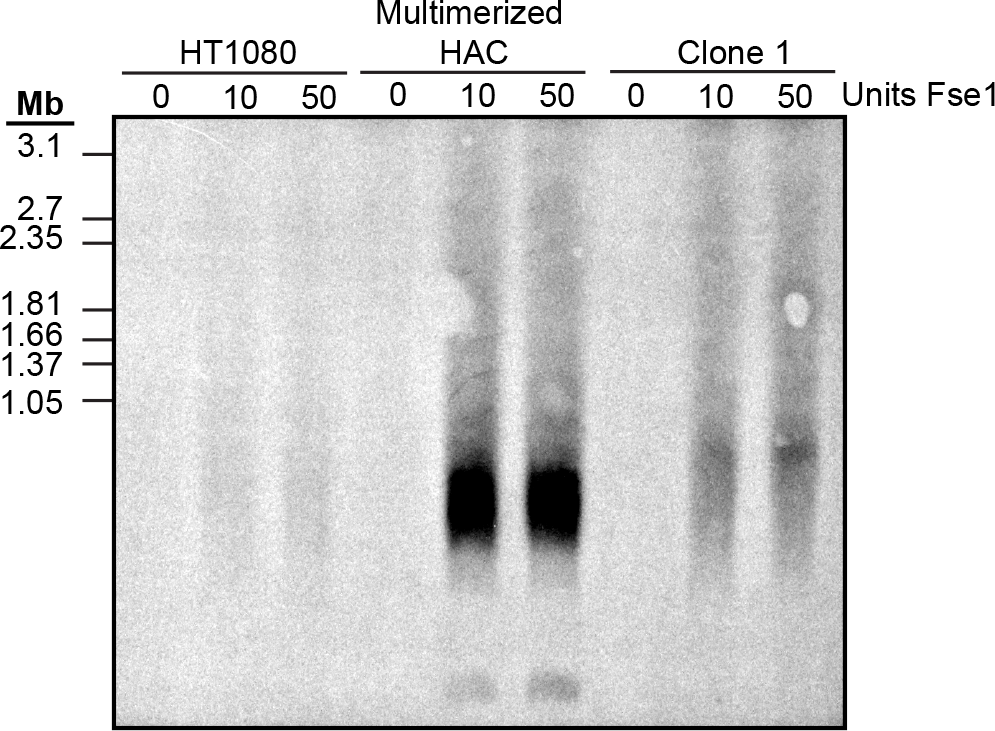
Southern blot analysis comparing HACs to the parental cell line as a control.

**Figure S6:**
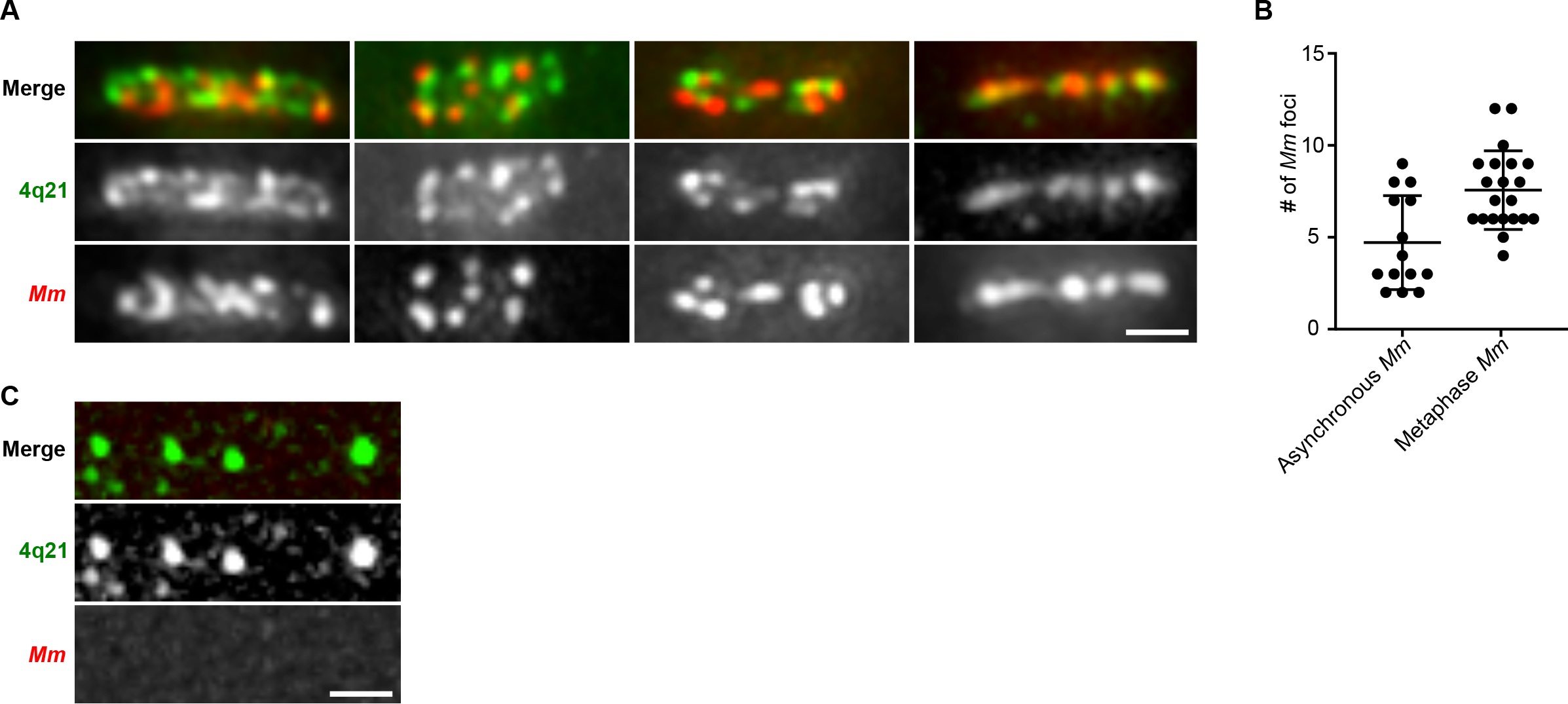
Representative examples of the HAC after physical stretching. A) 4q21 signal is typically more diffuse compared to *mycoides* signal suggesting that the two types of DNA have distinct chromatin properties. B) Quantification of *Mm* FISH foci in experiment shown in Fig. 4C,G. C) Representative images of stretched endogenous 4q21 labeled via FISH after enriching for cells in metaphase.

**Table S1:** Summary of the HAC clones generated in this study. The relevant figures that the HAC clones are described in is listed.

| Clone | Figures referenced | % of cells with HACs | Identifier |
| --- | --- | --- | --- |
| 1 | 2B,C; 4C-G; S5; S6 | 0.70 | yBB6 HAC 1-7 |
| 2 | 2A-E; 3B-D; 4A; S4 | 0.90 | yBB6 HAC 1-15 |
| 3 | 2B,C | 0.87 | yBB6 HAC 1-16 |
| 4 | 2B | 0.60 | yBB6 HAC 1-19 |
| 5 | 2B,C | 0.60 | yBB6 HAC 1-23 |
| 6 | 2B | 0.65 | yBB6 HAC 4-1-1 |
| 7 | 2B | 0.65 | yBB6 HAC 4-2-1 |
| 8 | 2B | 0.67 | yBB6 HAC 4-3-4 |

